# Characterization of the interaction domains between the phosphoprotein and the nucleocapsid of human Metapneumovirus

**DOI:** 10.1101/2021.06.02.446795

**Authors:** Hortense Decool, Benjamin Bardiaux, Luis Checa Ruano, Olivier Sperandio, Jenna Fix, Irina Gutsche, Charles-Adrien Richard, Monika Bajorek, Jean-François Eléouët, Marie Galloux

## Abstract

Human metapneumovirus (HMPV) causes severe respiratory diseases in young children. The HMPV RNA genome is encapsidated by the viral nucleoprotein, forming an RNA-N complex (N^Nuc^), which serves as template for genome replication and mRNA transcription by the RNA-dependent RNA polymerase (RdRp). The RdRp is formed by the association of the large polymerase subunit (L), which has RNA polymerase, capping and methyltransferase activities, and the tetrameric phosphoprotein (P). P plays a central role in the RdRp complex by binding to N^Nuc^ and L, allowing the attachment of the L polymerase to the nucleocapsid template. During infection these proteins concentrate in cytoplasmic inclusion bodies (IBs) where viral RNA synthesis occurs. By analogy to the closely related pneumovirus respiratory syncytial virus (RSV), it is likely that the formation of IBs depends on the interaction between P and N^Nuc^. However, the HMPV P-N^Nuc^ interaction still remains to characterize. Here, we finely characterized the binding domains involved in HMPV P and N^Nuc^ interaction by studying binding between recombinant proteins, combined with the use of a functional assay of the polymerase complex activity and the study of the recruitment of these proteins to IBs by immunofluorescence. We show that the last 6 C-terminal residues of HMPV P are necessary and sufficient for binding to N^Nuc^, that P binds the N-terminal domain of N (N_NTD_), and identified conserved N residues critical for the interaction. Our results allowed to propose a structural model of the HMPV P-N^Nuc^ interaction.

**IMPORTANCE:** Like respiratory syncytial virus (RSV), human metapneumovirus (HMPV) is a leading cause of severe respiratory infections in children but also affects human populations of all ages worldwide. Nowadays, no vaccine or efficient antiviral treatments are available for these two pneumoviruses. A better understanding of the molecular mechanisms involved in viral replication could help the design or discovery of specific antiviral compounds. In this work we have investigated the interaction between two major viral proteins involved in HMPV RNA synthesis, the N and P proteins. We finely characterized their domains of interaction, in particular a pocket on the surface of the N protein that could be used as a template for the design of compounds interfering with N-P complexes and viral replication.

## INTRODUCTION

Pneumonia is the leading cause of death among children younger than 5 years worldwide, and severe pneumonia is more frequently caused by viruses than bacteria (1). After the closely related respiratory syncytial virus (RSV), human metapneumovirus (HMPV) is recognized to be one of the most important cause of viral bronchiolitis and pneumonia in young children, causing 7 to 19% of all cases of acute respiratory tract infections (1, 2). HMPV infects mainly newborn, children, elderly and immunocompromised individuals worldwide. This virus was first reported in 2001 from Dutch children with acute lower respiratory tract illness, and serological studies have revealed that virtually every child has been exposed to HMPV by the age of 5 years (3). The clinical features of HMPV infection display as mid-to-upper respiratory tract infection, and can be severe enough to cause life-threatening bronchiolitis and pneumonia. As yet, there is no effective treatment or licensed vaccine for HMPV. Together with RSV, HMPV belongs to the *Pneumoviridae* family in the order *Mononegavirales* (4). HMPV is an enveloped virus that forms pleomorphic or filamentous virions. The virus genome is composed of a negative-sense single-stranded RNA of approximately 13.3 kb in size which encodes eight genes in the following order: 3’-N-P-M-F-M2-SH-G-L-5’ (5, 6). The M2 gene of HMPV contains two overlapping open reading frames (ORFs), encoding for M2-1 and M2-2 proteins which precise functions during HMPV replication remain unclear.

The HMPV genome is encapsidated by multiple copies of the nucleoprotein (N) forming helical nucleocapsids (N^Nuc^). This N^Nuc^ serves as template for genome replication and mRNA transcription by the viral polymerase complex formed by the large polymerase subunit (L) and its main cofactor the phosphoprotein (P). After virus binding to the cell surface and virus-cell membrane fusion, mediated by surface glycoproteins (F and G), nucleocapsids are released into the cytoplasm. Replication and transcription of the viral genome take place within viro-induced cytoplasmic inclusions named inclusion bodies (IBs) (7). These structures can be observed upon expression of P and N proteins alone (8), and it was recently shown for RSV that the interaction between P and N^Nuc^ is critical for the formation of IBs (9).

Among the components of the polymerase complex, P plays a pivotal role as a cofactor of the L polymerase but also as a molecular hub between viral and cellular partners. HMPV P, 294 amino acid residues in length, forms homo-tetramers. The atomic structure of the coiled-coil oligomerization domain (residues 171-194) was resolved by crystallography (10). Small angle X-ray scattering (SAXS) studies indicated that the flanking N-terminal (residues 1-170) and C-terminal (residues 195-294) regions (named PNT and PCT respectively) are intrinsically disordered, some of them, such as residues 195-237, having α-helical propensity (10, 11). More recently, the structure of the LP complex was resolved by cryo-EM (12). It revealed a tentacular arrangement of P, with each of the four protomers adopting a distinct conformation, demonstrating a “folding-upon-partner-binding” mechanism. Depending on the protomer, the L-binding region involved regions 171-236, 172-217, 170-231 or 169-264. On the other hand, by binding to the nucleocapsid, P mediates the attachment of the L protein to the N^Nuc^ template for viral RNA synthesis.

By analogy to RSV (13), the C-terminal domain of HMPV P is also thought to be the N^Nuc^ binding domain, but this has not been shown yet. It is noteworthy that the C-terminal extremity of P supposed to bind to N^Nuc^ was not visible in the crystal structure of the LP complex, indicating that this region is disordered in the absence of N^Nuc^. Furthermore, the encapsidation of neosynthesized genome or antigenome necessitates a pool of monomeric, RNA-free N, termed N^0^, which is kept in an unassembled state through an interaction with P which plays the role of molecular chaperone, until delivery to the sites of viral RNA synthesis. The crystal structures of recombinant HMPV N protein (394 residues) expressed in *E. coli* and purified either as an RNA-free monomeric N in complex with the N-terminal residues of P, or as rings of 10 N protomers complexed to RNA were resolved (14). In both states, the structures show that N presents two globular domains (N_NTD_ and N_CTD_) separated by a flexible linker that forms the RNA groove, and N- and C-arms. In the oligomeric state, the N- and C-arms play a key role in the interaction between N protomers and oligomerization, the N-arm binding to the flank of the N*i*−1 protomer and the C-arm binding atop the N*i* +1 protomer (*i* corresponding to the middle subunit of three adjacent N protomers). In the N^0^ state, the important conformational changes consist in packing of the C-arm of N in the RNA groove, impairing RNA binding.

In this work, based on the structural data of HMPV P and N proteins and their strong homologies with RSV N and P proteins, we finely characterized the binding domains involved in HMPV P-N^Nuc^ interaction. By combining biochemical and functional cellular assays, coupled with a rational mutational approach, we identified residues of P and N critical for their interaction. Our data show that the last C-terminal residues of P bind to the N_NTD_. These results allowed to establish a structural model of the interaction which could be used for the rational design of antivirals targeting the N^Nuc^-P interaction of HMPV.

## RESULTS

### The HMPV PCT domain allows to purify N-RNA rings expressed in bacteria

By analogy to RSV, it is thought that the N^Nuc^ binding domain of HMPV P is located at its C-terminus. For RSV, it was shown that co-expression of the C-terminal disordered region of P (PCT, residues 161-241) fused to GST together with N in bacteria allows the purification of complexes formed by GST-PCT and ring shape N-RNA oligomers (13, 15). Similarly, when expressed alone in *E. coli*, HMPV N also forms decameric rings containing RNA (14). In a first attempt to characterize the N^Nuc^ binding site of HMPV P, we thus decided to co-express recombinant HMPV GST-PCT (residues 200-294) and N proteins (from HMPV CAN 97-83 strain) in *E. coli*. The purified complexes were analyzed by SDS-PAGE stained with Coomassie blue. As shown on figure.1A, N was co-purified with GST-PCT to > 95% homogeneity. The PCT was then separated from GST by thrombin cleavage and the solubilized complex was analyzed by size exclusion chromatography, following optical density (OD) at 220, 260 and 280 nm (Fig. 1A). The elution profile showed a major peak (P1) with a OD_260nm_/OD_280nm_ ratio > 1 and an apparent mass of ~ 500 kDa, in agreement with the expected size of N-RNA decamers (Fig. 1B). A second peak of lower intensity (P2), with an apparent mass of ~ 10 kDa was detected only at 220 nm. This peak should correspond to PCT which does not present aromatic residues and a higher mass weight than predicted due to the elongated shape of this fragment. The fractions of P1 peak were pooled; analysis by SDS-PAGE stained with Coomassie blue of the sample showed the presence of a unique band corresponding to N (Fig. 1B). The separation of PCT and N upon gel filtration reveals the relatively low affinity of monomeric PCT for N. We then further analyzed the sample collected from the P1 peak by combining dynamic light scattering (DLS) and electron microscopy (EM) approaches. The profile of DLS measurement confirmed the homogeneity of the sample, with a single peak at 18 nm (Fig 1C), in agreement with the size of N-rings observed by EM (Fig 1D).

**Figure 1:**
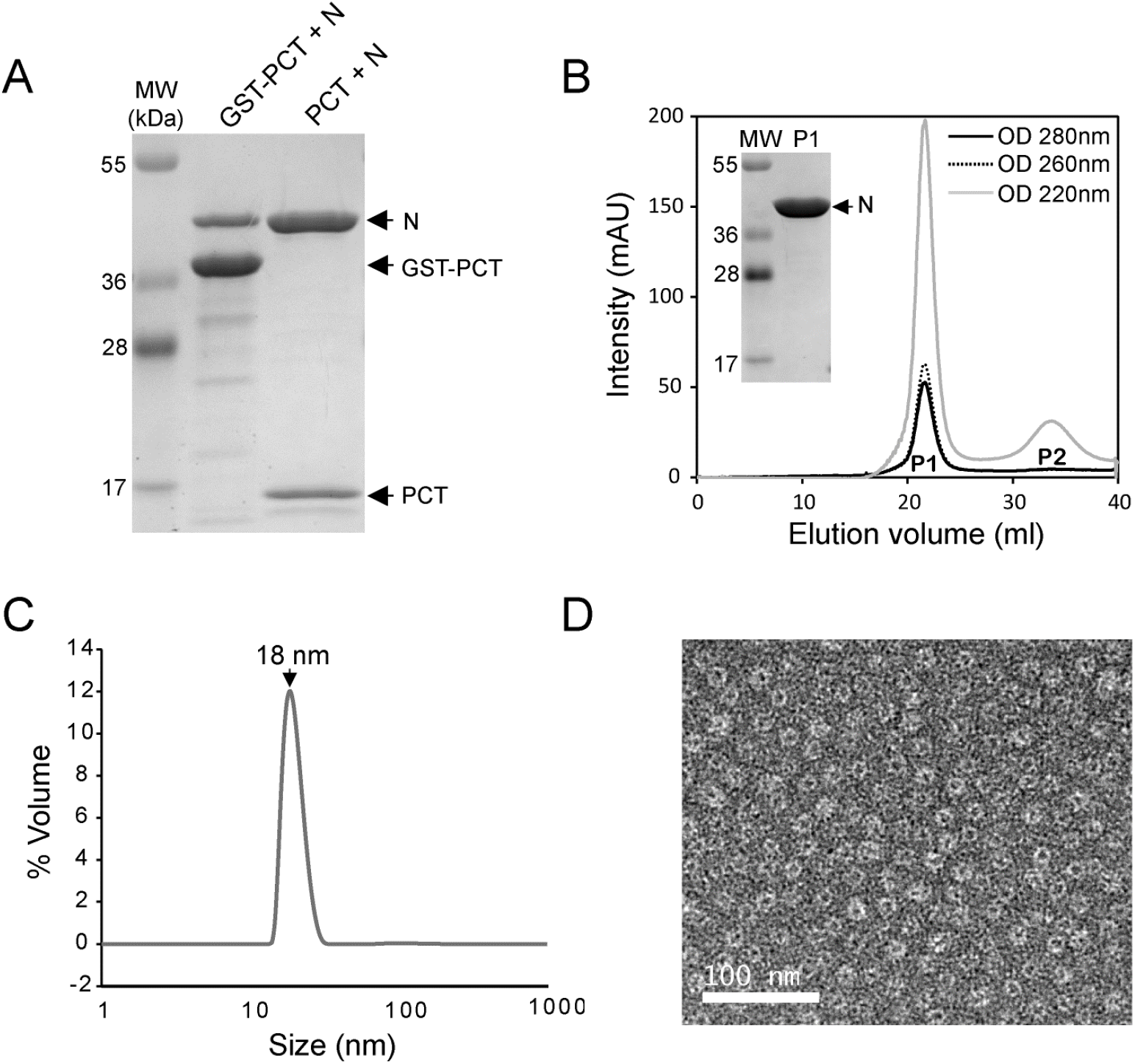
Purification of HMPV N-RNA rings using GST-PCT. A/ GST-PCT and N proteins were co-expressed in *E. coli*, followed by purification using the GST tag. The product of purification was analyzed by SDS-PAGE and Coomassie blue staining (left lane). The sample was then incubated in the presence of thrombin in order to cleave GST (remaining on beads) and isolate PCT-N in the supernatant (right lane), B/ Gel filtration profile of purified PCT-N complex. The curves corresponding to OD spectra at 220 nm, 260 nm, and 280 nm are presented. P1 and P2 indicate the two peaks detected. The fractions corresponding to P1 were pooled and the sample was analyzed by SDS-PAGE colored with Coomassie blue, C/ Dynamic light scattering (DLS) analysis of the purified N protein, showing a homogenous peak at 18 nm, corresponding to N oligomers. D/ Image of purified HMPV N-RNA rings as observed by negative stain electron microscopy. Bar, 100 nm.

Altogether, these results show that the GST-PCT fusion protein is sufficient to interact with nucleocapsid-like N-RNA rings.

### The last 6 residues of HMPV P constitute the minimum domain for N^Nuc^ binding

We thus investigated the minimal domain of HMPV P involved in N^Nuc^ binding. By analogy with RSV for which the 9 C-terminal residues of P are sufficient to interact with N (13), a series of GST fused peptides of 9 to 1 amino acid long derived from the C-terminal of P was generated. These constructs were co-expressed with N in *E.coli*, and their capacity to purify N was analyzed by SDS-PAGE and Coomassie blue staining. As shown on figure. 2A, the minimal sequence required for N purification corresponds to the 6 last C-terminal residues of P. In parallel, alanine scanning was performed with the GST-P[285-294] construct to characterize the residues of P involved in the interaction with N^Nuc^. Again, GST-P constructs and N were co-expressed in bacteria, followed by purification by Glutathione-Sepharose beads affinity, and co-purification of N was analyzed by SDS-PAGE and Coomassie blue staining. Only the four I289A, Y290A, L292A and M294A substitutions abrogated the interaction with N (Fig. 2B). In order to further investigate the potential role of residues Q291 and I293 of P in the interaction with N, the impact of the double mutations Q291A/I293A inserted into GST-PCT on N binding was also tested. These mutations did not affect N binding, confirming that these two residues are not directly involved in the interaction (Fig. 2C). These results confirm that the last residues of P are directly involved in the interaction with N. Of note, our results suggest that HMPV P-N interaction mainly involves hydrophobic interaction. As both hydrophobic and acidic residues of RSV P were previously shown to be critical for the interaction with N (Fig. 2D), these results suggest that the binding of HMPV P on N differs from RSV and involves specific interactions. We thus tested the capacity of each PCT fragments to pulldown N proteins. Figure 2E shows that HMPV PCT and RSV PCT did not allow to purify RSV N and HMPV N respectively. This last result confirms that HMPV and RSV N and P proteins cannot cross-interact, and that pneumoviruses P-N interactions are specific to each virus.

**Figure 2:**
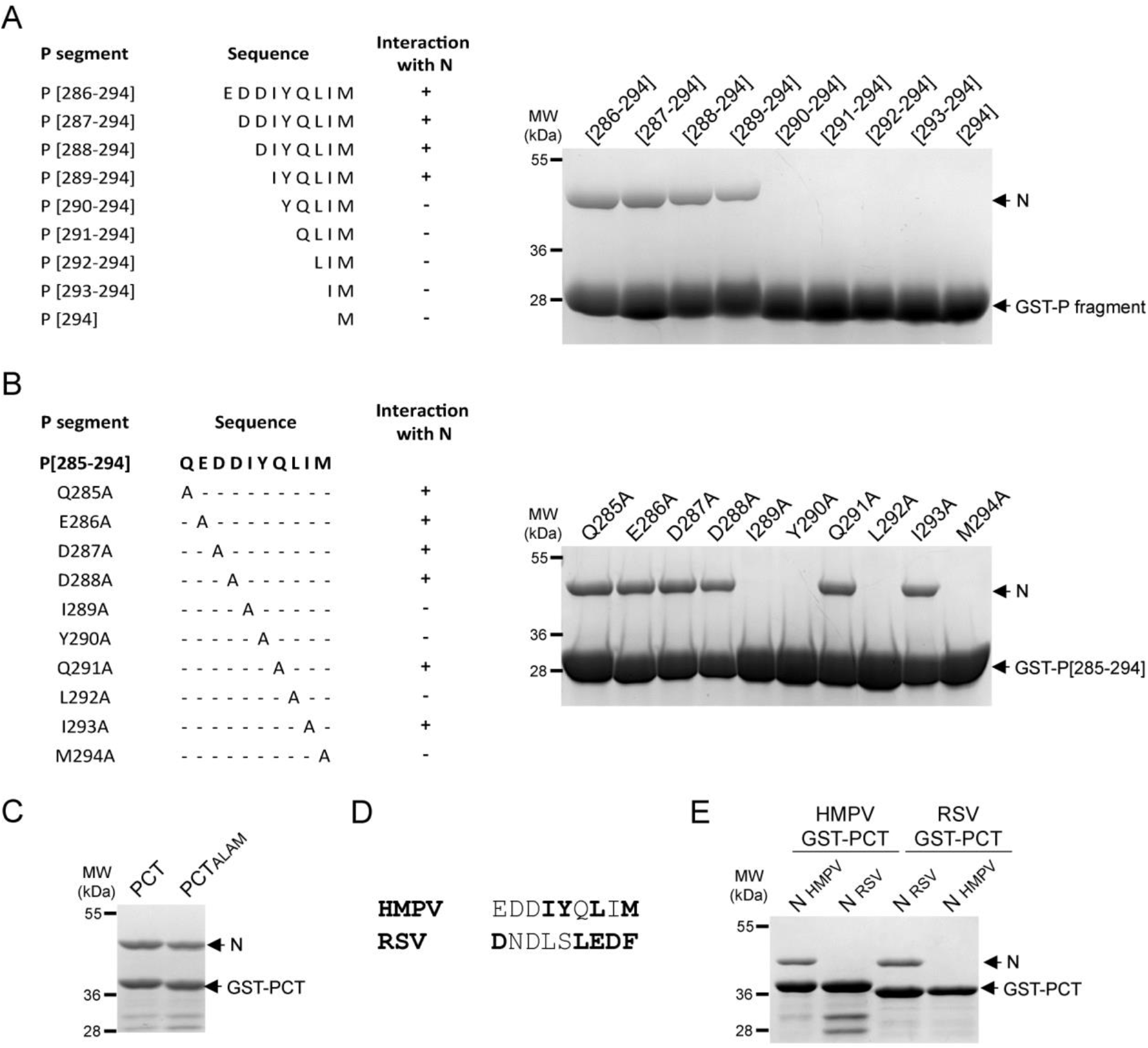
Identification of the residues of P involved in N binding. A/ GST-P fragments and B/ GST-P[285-294] (WT or mutants) (sequences indicated on the left) were co-expressed with N in bacteria, followed by purification using the GST tag. The products of purification were analyzed by SDS-PAGE and Coomassie blue staining (right). C/ GST-PCT or GST-PCT_ALAM_ corresponding to the double substitution of residues Q291 and I293 residues of P by alanine were co-expressed with N. The products of purification by GST were analyzed by SDS-PAGE and Coomassie blue staining. D/ Sequence alignment of the 9 last C-terminal residues of HMPV and RSV P proteins. The residues critical for the interaction with the N protein are indicated in bold. E/ HMPV and RSV GST-PCT (residues P(200-294) for HMPV and P(161-241) for RSV) constructs were co-expressed with either HMPV or RSV N proteins. Copurification of N proteins by GST-PCT fragment was analyzed by SDS-PAGE and Coomassie blue staining.

### Search for the P binding site on the surface of HMPV N

Although HMPV and RSV P-N^Nuc^ interactions have their own specificity and cannot cross-react, the strong homologies between N proteins suggest that the PCT binding site on N protein could be partially conserved. For RSV, the P binding site on N^Nuc^ is located at the surface of the N_NTD_. The residues of RSV N_NTD_ involved in the interaction with P, which are partially conserved between HMPV and RSV (Fig. 3A), were shown to form a well-defined pocket constituted of hydrophobic and positively charged residues (16, 17). Corresponding residues of HMPV N_NTD_ also form a pocket at the surface of the domain (Fig. 3B). In a first attempt to characterize the HMPV P binding domain on N, we thus tested the capacity of P to interact with the N_NTD_. The N_NTD_ was expressed in *E. coli* and purified using a 6xHis tag, showing that this domain is soluble when expressed alone (Fig. 3C). In parallel, the GST-P or GST-PΔM294 (deletion of the last C-terminal residue of P) proteins were expressed in bacteria either alone and purified using the GST tag, or co-expressed with N_NTD_ followed by purification by 6xHis tag. Analysis of the samples by SDS-PAGE showed that N_NTD_ allowed to co-purify the wild type P, and that deletion of the last residue of P was sufficient to impair the interaction (Fig. 3C). We then substituted by alanine the residues of the HMPV N_NTD_ identified by analogy with RSV, i.e. residues L46, L47, E50, Y53, D128, R132, M135, R151, P152, and S153 (Fig. 3B). Again, the capacity of GST-P to co-purify N mutant proteins was tested. Whereas mutants L46A, L47A, and P152 could still interact with P, only a weak band of N was detected for mutants E50A, M135A, R151A and S153A, and mutations Y53A, D128A, R132A abrogated the interaction (Fig. 3D). Of note, it was previously shown that similar punctual mutations did not impact RSV N folding and solubility, suggesting that these mutations on HMPV N only affect the capacity of P to interact with N. These results thus reveal a critical role of hydrophobic residues, but also of negatively and positively charged residues of N_NTD_ in P binding. Overall, these data confirm that the P binding site on N presents strong homologies between HMPV and RSV.

**Figure 3:**
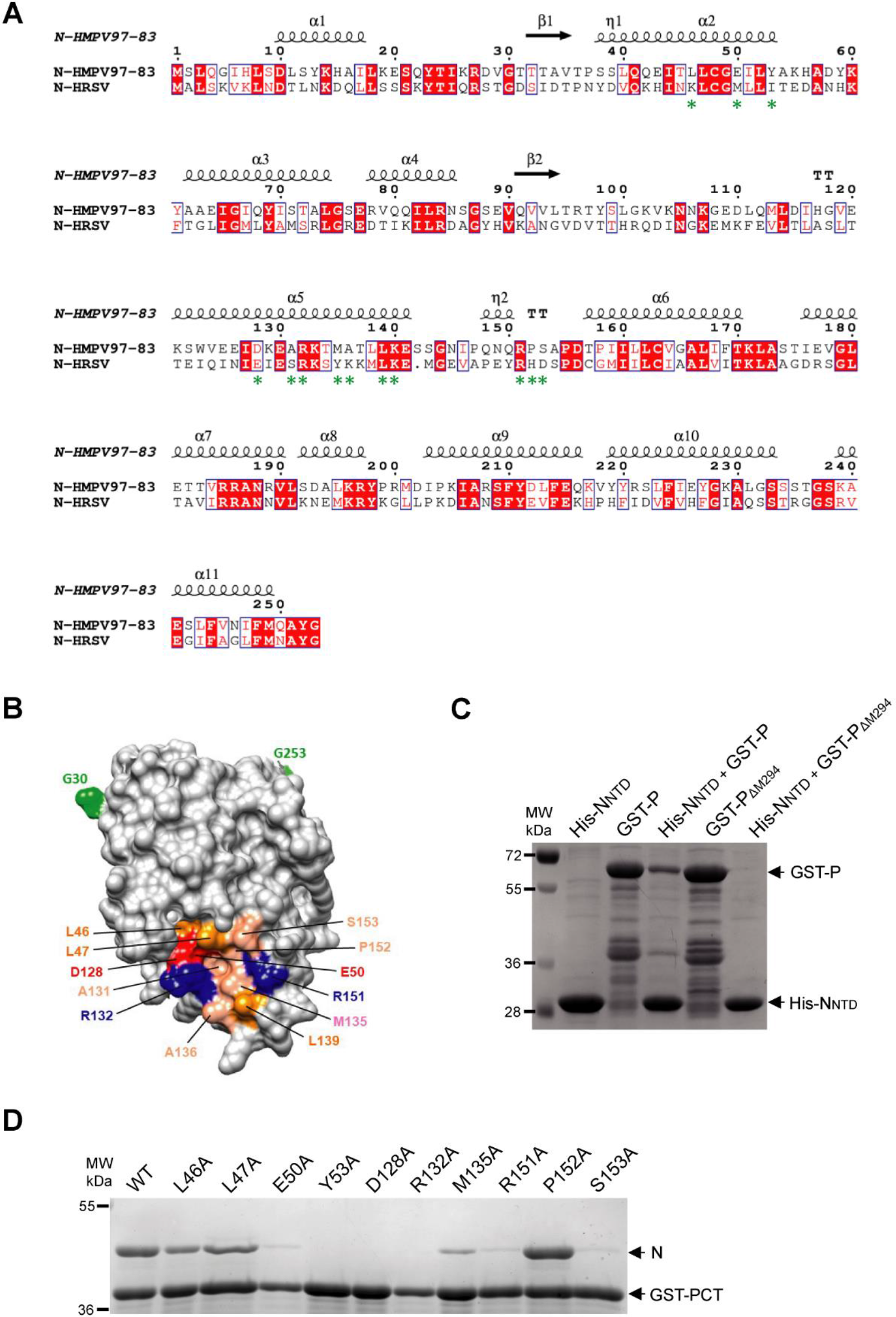
Search for the PCT binding site on HMPV N. A/ Amino acid sequence alignments between the N-terminal domains of N (residues 1-252/253) proteins of HMPV (strain CAN 97-83) and human RSV (HRSV, strain Long VR-26). Invariant residues are highlighted in white font on a red background. The secondary structure elements observed in the crystal structure of HMPV N protein (11) are indicated above the sequence. Green asterisks below the sequence indicate the residues constituting the P binding pocket of RSV N. Uniprot accession codes: NCAP_HRSVA (Human RSV); A1DZS3_9MONO (Human MPV). Sequences were aligned with Clustal W and treated with ESPript 3. B/ Surface representation of HMPV N_NTD_ (from Gly30 to Gly 253, indicated in green) showing the potential P binding pocket, with acidic amino acids colored in red, basic residues in blue, and hydrophobic residues in orange, C/ GST-P or GSP-P_ΔM294_ and N_NTD_-6xHis proteins were co-expressed in *E. coli*, followed by purification using the 6xHis tag. GST-P or GSP-P_ΔM294_ alone were purified using the GST-tag. The products of purification were analyzed by SDS-PAGE and Coomassie blue staining, D/ GST-PCT and mutant N proteins were co-expressed in *E. coli*, followed by purification using the GST tag. The product of purification was analyzed by SDS-PAGE and Coomassie blue staining.

### Validation of P-N^Nuc^ binding domains in eukaryotic cells

As P-N^Nuc^ interaction is required for viral polymerase activity, we then studied the impact of selected mutations of P (M294A) or N (R132A and R151A) on HMPV replication/transcription using a bicistronic subgenomic minigenome, pGaussia/Firefly. This construct contains the Gaussia and Firefly luciferase genes under the control of gene start and gene stop sequences, as well as Leader and Trailer sequences of HMPV genome (Fig. S1). Briefly, BSRT7 cells were transfected with plasmids coding for N (N_WT_, N_R132A_ or N_R151A_), P (P_WT_ or P_M294A_), L, and M2-1 proteins, the plasmid pGaussia/Firefly minigenome, and pSV-β-Gal (used to normalize transfection efficiencies). In this system, expression of luciferases depends on the HMPV polymerase complex activity which can thus be quantified by luminescence measurement. As shown on figure 4A, mutations M294A of P or R132A of N abrogated HMPV polymerase activity and R151A induced a strong decrease on its activity, with only 30% of activity compared to the control condition, although the levels of expression of these proteins were similar to those of wild type proteins (Fig. 4B). These results correlate with those of pulldown assays, the mutant N_R151A_ that still presented a weak interaction with P displaying a residual polymerase activity.

**Figure 4:**
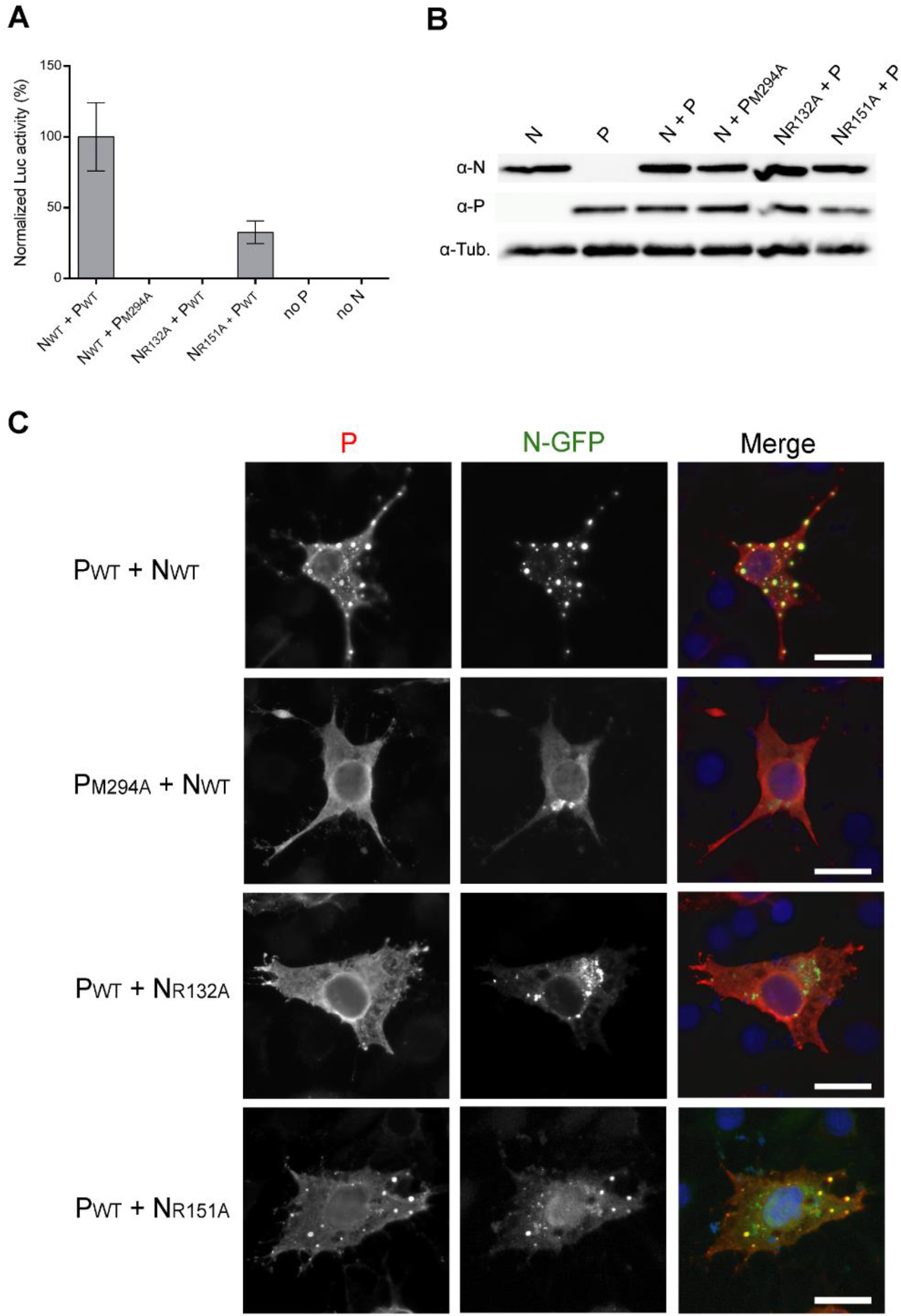
The residues involved in P-N^Nuc^ interaction are critical for the polymerase activity and the formation of inclusion bodies. A/ HMPV polymerase activity in the presence of P and N mutants. BSRT7/5 cells were transfected with plasmids encoding the P (WT or M294A mutant), N (WT, or R132A and R151A mutants), L, and M2-1 proteins, the pGaussia/Firefly minigenome, together with pCMV-βGal for transfection standardization. Viral RNA synthesis was quantified by measuring the Firefly luciferase activity after cell lysis 24 h after transfection. Each luciferase minigenome activity value was normalized based on β-galactosidase expression and is the average of three independent experiments performed in triplicate. Error bars represent standard deviations calculated based on three independent experiments made in triplicate. B/ Western blot showing the expression of N protein variants in BSRT7/5 cells. C/ Impact of P and N mutations on the formation of HMPV cytoplasmic IBs. N-GFP and P (or mutants) proteins were coexpressed in BSRT7/5 cells; cells were then fixed 24 h post-transfection, labeled with anti-P antibody and the distribution of viral proteins was observed by fluorescence microscopy. Nuclei were stained with Hoechst 33342. Scale bars, 10 μm.

In HMPV-infected cells, P-N^Nuc^ interaction induces the formation of inclusion bodies (8). For RSV, we have shown previously that IBs correspond to viral factories where viral RNA synthesis occurs (18). We then assessed the impact of these mutations on the capacity of N and P to form pseudo-IBs when co-expressed in cells. To facilitate N detection, N-GFP fusion protein was generated to allow its detection by epifluorescence. Based on previous studies on RSV P-N^Nuc^ interaction, the GFP was fused to the C-terminus of N and this construct was co-expressed with wild-type N at a ratio of 1:1. Cells were thus co-transfected with plasmids encoding N-GFP, N (both WT or mutants R132A or R151A), and P (WT or mutant M294A). The P protein was detected by immunolabelling using rabbit anti-P antibody, and nuclei were stained with Hoechst 33342. When expressed together in BSRT7 cells, WT N-GFP/N and P were found to co-localize into cytoplasmic inclusions similar to IBs observed upon HMPV infection (7, 8) (Fig. 4C). On the contrary, when expressing P_M294A_ in the presence of N_WT_ or P_WT_ in the presence of N_R132A_, P presented a diffuse cytoplasmic distribution and N protein was mostly detected in cytoplasmic aggregates (Fig. 4C). These observations suggest that in the absence of N-P interaction, overexpressed N presents a strong tendency to aggregate. Finally, co-expression of the mutant N_R151A_ with P still allowed to observe IBs, although less numerous and smaller than those observed in the presence of WT proteins. This last result correlates with previous results showing that mutation R151A of N drastically reduced but did not abrogate N-P interaction and polymerase activity. Altogether, these results confirm that the residues of P and N previously identified as critical for N^Nuc^-P interaction *in vitro* are also critical for the formation of IBs and polymerase activity in cells.

### Molecular modelling of PCT-N_NTD_ interaction

To gain insight into the possible binding mode of P with N_NTD_, we built models of the last 6 residues of P (peptide I289-YQLI-M294 named P6) bound to N, by analogy with the PCT-N_NTD_ complex of RSV. For RSV, only the last 2 residues of P (D240-F241) were resolved in the PCT-N_NTD_ X-ray structures (17), and are bound to a hydrophobic pocket of N similar to the one we characterized at the surface of HMPV N. Using comparative modeling and refinement with HADDOCK (19), we obtained three clusters of HMPV P6/N_NTD_ complex structures representing possible binding modes of P6 (Table S1). The first two clusters in terms of HADDOCK-score were the most populated and displayed similar Root-mean-square deviation (RMSD) among their members (considering only the last three residues of P, i.e. residues L292-I293-M294, called P3). In those two clusters, the C-terminal carboxyl group of M294 is involved in an ionic interaction with the guanidinium of either R131 or R152 side-chains of N. Additionally, only L292 and M294 of P3 consistently display hydrophobic contacts with N. This result is consistent with our mutation data showing that I293 of P is not required for binding to N, contrary to L292 and M294. The 10 best-scoring structures from cluster 1 and 2 are shown in Fig. S2, superimposed with the RSV P2/N_NTD_ structure (residues D240-F241 of P). Owing to the best HADDOCK-score and non-bonded energies of cluster 2, we considered this cluster as most representative for the potential binding mode of HMPV P3 to N_NTD_. Interestingly, more than half of the structures from this cluster display a hydrogen-bond between the backbone oxygen of I293 of P3 with the side-chain of R151 of N, while the side-chain of R132 of N forms a salt-bridge with the C-terminal carboxyl group of M294 of P. Thus, the interactions determined by molecular simulation correlate with the experimental data showing the importance of R132 and R151 of N and of M294 of P for PCT-N interaction.

The interactability propensity of each modeled N_NTD_ conformation was further profiled with our in-house tool InDeep in order to select a subset of N_NTD_ models being more likely to interact with the P3 peptide. On this subset of N_NTD_ conformations, the hydrophobic channel of InDeep was also used to locate regions where hydrophobic moieties of P3 are expected to interact at the N_NTD_ surface. Three hydrophobic patches are detected on the N_NTD_ surface, that present a good match with hydrophobic moieties of P3 structures belonging to cluster 2 of HADDOCK: i) one occupied by L292 of P3; ii) another occupied by M294 of P3; and iii) the last one located in an unoccupied region of the N pocket (Fig. 5A). Based on experimental and modelling information, the structure shown in Figure 5 represents a consistent model of P3/N_NTD_ complex. The N_NTD_ conformation of this structure is predicted to be favorable to protein-protein interaction by InDeep, and the P3 conformation, belonging to cluster 2 of HADDOCK, binds to N_NTD_ with a combined effect of electrostatic/polar interactions and hydrophobic contacts in agreement with the mutational experiments. In addition, a procedure of virtual alanine scanning using the same InDeep approach confirmed some of the mutagenesis results, such as the importance of R132, M135, and R151 of N for the interaction with P. It also suggests that residue L139 of N may be key for the N_NTD_ patch for partners’ binding.

**Figure 5:**
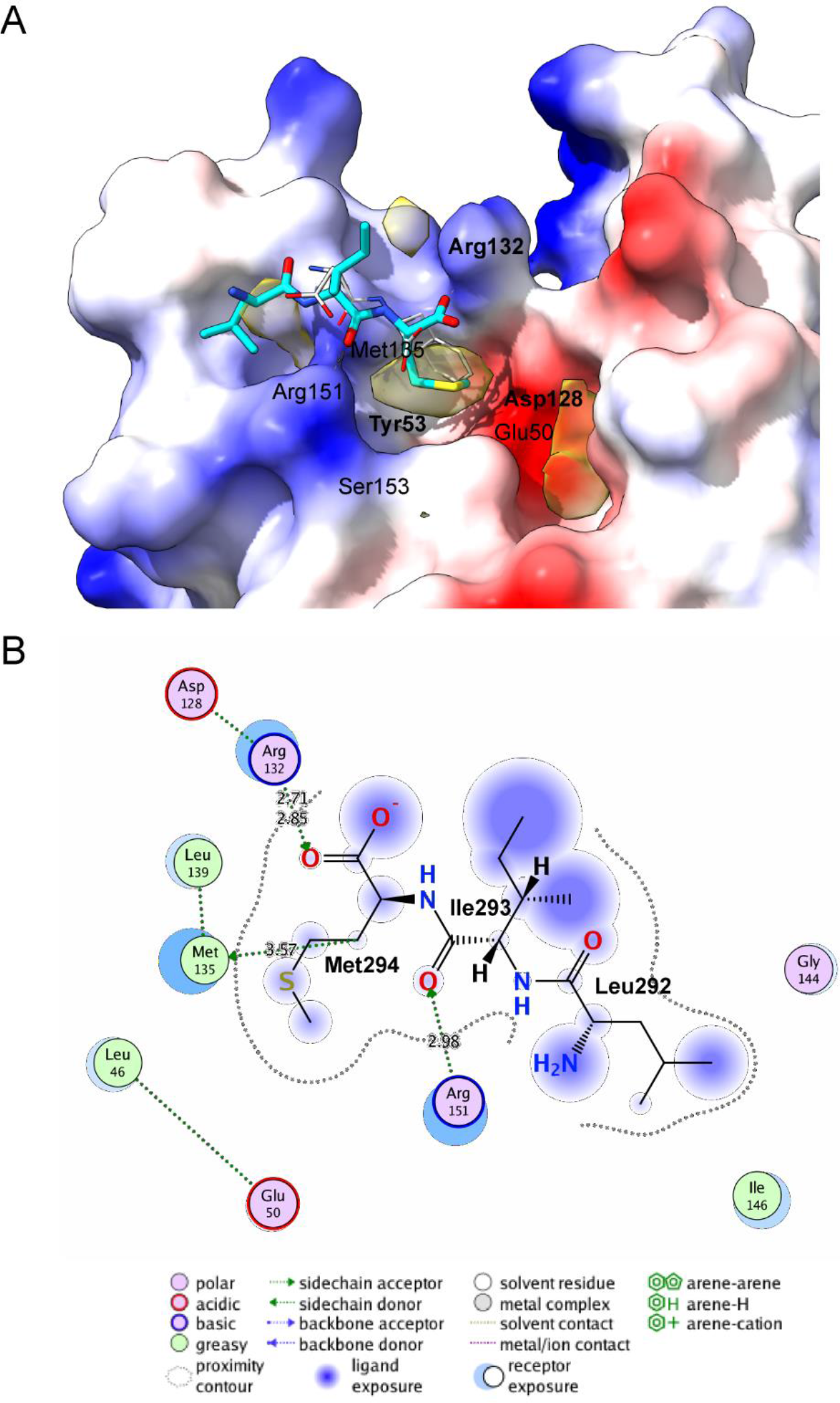
Model of HMPV P3/N_NTD_ complex from HADOCK refinement. A/ The N_NTD_ protein is colored by electrostatic surface and the P3 peptide (residues L292-I293-M294 of P) is shown as sticks. The residues of N_NTD_ critical for the interaction with P are indicated, with residues for which mutation to alanine abrogate P binding are labeled in bold. Yellow Surface shows InDeep hydrophobic channel prediction where hydrophobic contacts are expected to occur at the N_NTD_ surface. B/ Interaction diagram between P3 peptide (Leu292-Ile293-Met294) and N_NTD_. Legend indicated below the diagram.

These analyses reveal that despite the different physico-chemical nature of residues involved in the interaction, PCT binding at the N surface is structurally similar between HMPV and RSV.

## DISCUSSION

Beyond its interest in better understanding the molecular mechanisms of virus replication, the study of protein-protein interactions required for Pneumoviruses’ polymerase complex activity also presents interest for the development of specific antiviral compounds. The activity of this complex depends on multiple highly specific protein-protein interactions that have no cellular counterparts. Among those, N^Nuc^-P interaction which is critical for viral polymerase activity constitutes a singular potential target for antivirals. We have previously described RSV N^Nuc^-P interaction at the molecular level, showing that the C-terminus of P binds to a well-defined pocket at the surface of N (9, 13, 16). We also made the proof of concept that small chemical compounds can target this pocket of N and inhibit viral replication in cells (17). More recently, Sá et al. showed that flavonoids derived molecules could also specifically bind to N pocket and block its interaction with P (20). Given the strong homologies between RSV and HMPV N and P proteins, it could be expected that a similar strategy could be used to block HMPV replication. It is noteworthy that ideally, the discovery of compounds that could inhibit replication of these two closely related viruses at the same time would be much more advantageous.

In the present study, we investigated the domains involved in HMPV P-N^Nuc^ interaction using pulldown assays between recombinant proteins (either truncated or mutated proteins) expressed in *E. coli*. As for RSV, the HMPV N protein was purified as nanorings formed by N and bacterial RNA (Fig 1). Our results revealed that mechanisms of binding between P and N^Nuc^ are similar between RSV and HMPV, the C-terminus of P interacting with a pocket located on the surface of the N N-terminal globular domain N_NTD_. However, it appeared that P and N amino acid residues involved in these interactions have different contributions/roles between RSV and HMPV, and that the P-N^Nuc^ interaction is specific of each virus. More specifically, we showed that the minimal domain of HMPV P required for N binding is constituted by the last 6 C-terminal residues of P which are mostly hydrophobic residues (IYQLIM), and determined that only Q291 and I293 are not critical for the interaction (Fig. 2). This P fragment is thus shorter than for RSV for which N binding minimal domain involves the 9 C-terminal residues of P and is composed of both hydrophobic and acidic residues (13, 16). We also identified 7 residues of N involved in P binding, *i.e*. residues E50, Y53, D128, R132, M135A, R151A and S153A, which form a well-defined pocket at the N_NTD_ surface (Fig 3). Of note, the pivotal role of residues R132 and R151 of N for polymerase activity and IBs formation sustained *in vitro* binding assays (Fig. 4). Of most interest, residues R132 and R150 of RSV N were also shown to play a key role in P-N^Nuc^ interaction, the aliphatic part of the R132 side chain being involved in the stacking of the aromatic ring of the residue F241 of P. In order to better understand the binding mode of HMPV P to N, molecular modelling was thus performed and the best structural models support that R132 and R151 of N and of M294 of P are critical for PCT-N interaction (Fig. 5). Nevertheless, molecular modelling was restricted to the binding of the last 3 C-terminal residues of P, and did not allow to predict the interaction of all the 6 critical residues of P on N. As also observed in the crystal structures of RSV PCT/N_NTD_ complexes (17), the last C-terminal residue of P seems to drive the binding to a well-defined N_NTD_ pocket and this primary binding could allow transient contacts with upstream P residues outside of the N pocket.

In conclusion, our data confirm the strong structural homologies between HMPV and RSV P-N complexes but also highlight some singularities. These data suggest that the HMPV P binding pocket at the N_NTD_ surface could represent a new target for the rational design of antivirals.

## MATERIALS AND METHODS

### Plasmid constructions

The N and P sequences used for bacterial expression were derived from HMPV CAN97-83 strain. The sequence of N or N_NTD_ (residues 30-255 of N) were cloned into pET-28a(+) vector (Novagen) at NcoI/XhoI or BamHI/XhoI restriction sites respectively, to allow bacterial expression of full-length N or N_NTD_ domain with a C-terminal or N-terminal 6xHis tag. The sequence of P or PCT (residues 200-294 of P), were cloned into pGEX-4T3 vector, at BamHI and SmaI sites to express in bacteria GST-P and GST-PCT fusion proteins. For the construction GST-P[285-294], complementary antiparallel oligonucleotides were hybridized and inserted at the BamHI/SmaI sites in the pGEX-4T-3 vector.

A HMPV minigenome plasmid containing Gaussia/Firefly luciferases was designed and synthesized by Genscript, cloned in pUC57 vector, and containing the trailer, leader, gene start (GS) and gene end (GE) sequences derived from HMPV CAN97-83 strain (see Fig S1). The first ORF of this pGaussia/Firefly minigenome codes for the Firefly luciferase and the second one codes for the Gaussia luciferase. The plasmids pP, pN, pL, and pM2-1 corresponding to the sequences of NL99-1 HMPV strain cloned into pCite vector. For expression of N-GFP fusion protein, the mGFP gene was amplified by PCR and cloned in frame at the 3’ end of N at the EcoRI site of pN, using In-fusion HD cloning kit (Takara Bio). Point mutations were introduced in the P and N sequences by site-directed mutagenesis using the Q5 site-directed mutagenesis Kit (New England Biolabs). Sequence analysis was carried out to check the integrity of all constructs. All the oligonucleotides sequences are available on request.

### Antibodies

Antisera used in this study included polyclonal rabbit antisera raised against recombinant HMPV N and P expressed in bacteria. A mouse monoclonal anti-β-tubulin (Sigma) and secondary antibodies directed against mouse and rabbit Ig G coupled to HRP (P.A.R.I.S) were also used for immunoblotting. Secondary antibody directed against rabbit Ig G coupled to Alexafluor-594 (Invitrogen) was used for immunofluorescence experiments.

### Cell culture and transfections

BHK-21 cells (clone BSRT7/5) constitutively expressing the T7 RNA polymerase (21) were grown in Dulbeco Modified Essential Medium (Lonza) supplemented with 10% fetal calf serum (FCS), 2 mM glutamine, and antibiotics. The cells were grown at 37°C in 5% CO2, and transfected using Lipofectamine 2000 (Invitrogen) as described by the manufacturer.

### Minigenome replication assay

BSRT7/5 cells at 90% confluence in 48-well dishes were transfected using Lipofectamine 2000 (Invitrogen) with a plasmid mixture containing 0.125 μg of pGaussia/Firefly minigenome, 0.125 μg of pN, 0.125 μg of pP (WT and mutants), 0.06 μg of pL, 0.03 μg of pM2-1, as well as 0.03 μg of pSV-β-Gal (Promega) to normalize transfection efficiencies. Cells were harvested at 24 h post-transfection and lysed in Firefly lysis buffer (30 mM Tris [pH 7.9], 10 mM MgCl2, 1 mM dithiothreitol [DTT], 1% [vol/vol] Triton X-100, and 15% [vol/vol] glycerol). The Firefly luciferase activity was determined for each cell lysate with an Infinite 200 Pro (Tecan, Männedorf, Switzerland) and normalized based on β-galactosidase (β-Gal) expression. Transfections were done in triplicate, and each independent transfection experiment was performed three times. For proteins expression analysis, cells were lysed in Laemmli buffer and analyzed by Western blotting (WB) using anti-N, anti-P, and anti-tubulin antibodies according to standard protocols.

### Fluorescence microscopy

Immunofluorescence microscopy was performed with cells grown on coverslips and previously transfected with pN-GFP, pN (both WT or mutants R132A and R151A, at a ratio 1:1), and pP (WT or mutant M294A). At 24 h post transfection, cells were fixed with 4% paraformaldehyde (PFA) for 25 min, made permeable, and blocked for 30 min with PBS containing 0.1% Triton X-100 and 3% bovine serum albumin (BSA). Cells were then successively incubated for 1 h at room temperature with primary and secondary antibody mixtures diluted in PBS containing 0.3% BSA. For nucleus labeling, cells were exposed to Hoechst 33342 stain (Invitrogen) during incubation with secondary antibodies. Coverslips were mounted with ProLong Gold antifade reagent (Invitrogen) and observed with an inverted fluorescence microscope (Zeiss Axiovision). Images were processed with ZEN software (Zeiss) and ImageJ software.

### Expression and purification of recombinant proteins

*E. coli* BL21 bacteria (DE3) (Novagen) transformed with pGEX-P plasmids (WT or mutant) or pET-N_NTD_ were grown at 37°C for 8 hours in 100 ml of Luria Bertani (LB) medium containing 100μg/ml ampicillin or 50μg/ml kanamycin, respectively. Bacteria transformed with pET-N-derived plasmids together with pGEX-P derived plasmids were grown in LB medium containing ampicillin (100 μg/ml) and kanamycin (50 μg/ml). The same volume of LB was then added and protein expression was induced by adding 80μg/ml isopropyl-ß-D-thio-galactoside (IPTG) to the medium. The bacteria were incubated for 15 hours at 28°C and then harvested by centrifugation. For purification using the GST-tag, bacteria were re-suspended in lysis buffer (50 mM Tris-HCl pH 7.8, 60 mM NaCl, 1 mM EDTA, 2 mM DTT, 0.2% Triton X-100, 1 mg/ml lysozyme) supplemented with complete protease inhibitor cocktail (Roche), incubated for 1 hour on ice, sonicated, and centrifuged at 4°C for 30 min at 10,000g. Glutathione-Sepharose 4B beads (GE Healthcare) were added to clarified supernatants and incubated at 4°C for 3 hours. Beads were then washed two times in lysis buffer and three times in PBS 1X, then stored at 4°C in an equal volume of PBS. To isolate GST-free P fragments or N rings, beads containing bound proteins were incubated with thrombin (Novagen) overnight at 20°C. Purified recombinant N proteins were loaded onto a Superdex 200 16/30 column (GE Healthcare) and eluted in 20 mM Tris/HCl pH 8.5, 150 mM NaCl. For purification of N_NTD_-6xHis fusion protein purification, bacterial pellets were re-suspended in lysis buffer (20 mM Tris-HCl pH8, 500 mM NaCl, 0.1% TritonX-100, 10 mM imidazole, 1 mg/ml lysozyme) supplemented with complete protease inhibitor cocktail (Roche). After sonication and centrifugation, lysates were incubated 30 min with chelating Sepharose Fast Flow beads charged with Ni^2+^ (GE Healthcare). Beads were then successively washed in the washing buffer (20 mM Tris-HCl, pH 8, 500 mM NaCl) containing increasing concentration of imidazole (25, 50, and 100 mM), and proteins were eluted in the same buffer with 500 mM imidazole. For SDS-PAGE analysis, samples were prepared in Laemmli buffer, denatured 5 min at 95 °C, separated on 10% plolyacrylamide gel and detected by Coomassie brilliant blue.

### Dynamic light scattering (DLS)

Size measurement of purified N oligomers was performed at 20 °C using a helium-neon laser wavelength of 633 nm and detection angle of 173° with a Zetasizer Nano (Malvern). Ten measurements were made, with an acquisition time of 10 s for each measurement. Hydrodynamic diameters (*DH*) were calculated using the DLS software provided by the instrument manufacturer. The results were presented as size distribution (nm).

### Negative stain electron microscopy observations of recombinant nucleoproteins

Three microliters of sample were applied to the clean side of carbon on a carbon–mica interface and stained with 2% sodium silicotungstate. Micrographs were recorded on a FEI Tecnai T12 microscope operated at 120 kV with a Gatan Orius 1000 camera. Images were recorded at a nominal magnification of 23 000 × resulting in a pixel size of 2.8 Å.

### Molecular modelling

In the X-ray structures of HMPV N^Nuc^ (PDB 5fvc) or N^0^-P (PDB 5fvd), the N_NTD_ residues L110 to M113, corresponding to a long beta-hairpin close to the PCT binding site, are not modeled. We used Modeller (22) to first construct a complete model HMPV N_NTD_ (residues 32 to 251) using as templates the X-ray structures of HMPV N^Nuc^ (residues 32 to 109 and 114 to 251) and RSV N_NTD_ (PDB 4uc6) for the missing beta-hairpin (residues 110 to 113). Next, using Modeller again, we generated 100 models of HMPV N_NTD_ complexed with the last six residues of P (peptide I289-YQLI-M294 called P6) using the RSV N_NTD_ structure in complex with the last two residues of RSV P (PDB 4uc9), *i.e.* D240-F241 (called P2), as template for HMPV P6 by aligning the last 2 residues of HMPV-P and RSV-P. To compensate for possible alternate conformations of exposed residues in apo and holo-form of N_NTD_, no template was used for R132 and R151 side-chains. Next, P6-N_NTD_ models where M294 side-chain of P is pointing towards the N_NTD_ hydrophobic pocket (by analogy with the binding of F241 of P on N RSV) were selected, to obtain an ensemble 41 P6-N_NTD_ models. Finally, the models were submitted to a water refinement procedure using the refinement interface of the HADDOCK2.2 server (23). Each input P6-N_NTD_ complex model was refined 20 times to obtain 820 refined models. The best 400 models with the lowest HADDOCK-score were clustered based on similarity of the L292-I293-M294 residues of P, called P3 (I289-Y290-Q291 were not considered for clustering since they appear very floppy in the refined models and had no structural templates during homology modeling). Models were first superposed on their N_NTD_ backbone atoms and the RMSD was computed on the backbone atoms of L292-I293-M294 of P. Clustering was performed using the HADDOCK tool *cluster_struc* (24) with a 2Å cut-off and a minimal cluster size of 10.

In addition, the 820 P6-N_NTD_ conformations generated by HADDOCK were profiled using InDeep software in order to select the most suitable N_NTD_ conformations to bind P3. InDeep is a deep-learning model based on a FCN (Fully Convolutional Network) trained from known structures of protein complexes capable of predicting protein-protein interaction interfaces. Each N_NTD_ pocket conformation is placed within a 3D grid composed of 1 Å^3^ voxels. The model computes a probability value for each voxel representing its interaction propensity. The predictions were performed on the putative P6 binding site on N_NTD_ surface. An “interactability-score” of the pocket is computed by taking the mean of the 150 best voxels probabilities. This value represents the binding propensity of a given N_NTD_ pocket conformation. Then, on the 200-top conformations with the highest “interactability-score” the hydrophobic channel of InDeep was applied in order to locate the most probable hydrophobic patches on the N_NTD_ pocket where P3 hydrophobic side chains could interact. In addition, once a set most promising conformations was identified, a virtual alanine scanning was performed using the same tool but by mutating one by one the residues surrounding the patch of interaction of P3. The results are provided as a set of correlations, one for each residue, highlighting the impact of the simulated mutation on the interactability of the N_NTD_.

## ACKNOWLEDGMENTS

We thank the immunologist’s team of Seppic laboratory (Maisons-Alfort, France) and the animal facilities of Anses, Maisons-Alfort for rabbit immunization and collect of anti-N and anti-P antisera, P. Collins (NIH, Boston) and S. Biacchesi (INRAE, Jouy-en-Josas) for the sequences of HMPV CAN97-83 N and P genes, B. van den Hoogen (Erasmus MC, Rotterdam) for the plasmids of HMPV NL99-1 strain. This work was carried out with the financial support of the French Agence Nationale de la Recherche, specific program ANR Decrisp n° ANR-19-CE11-0017.

We declare that we have no conflicts of interest with the contents of this article. H.D., M.G. and J.-F.E. designed experiments. H.D., M.B. performed molecular, and cellular assays. B.B., L.C.R. and O.S. performed molecular simulations. I.G performed electron microscopy observations. C.-A.R. performed gel filtration and *in vitro* study. M.G. wrote the paper with contributions from all authors, and J.-F.E. and M.G. edited the manuscript. All authors commented on the manuscript.

---

**Supplementary Figure 1:**
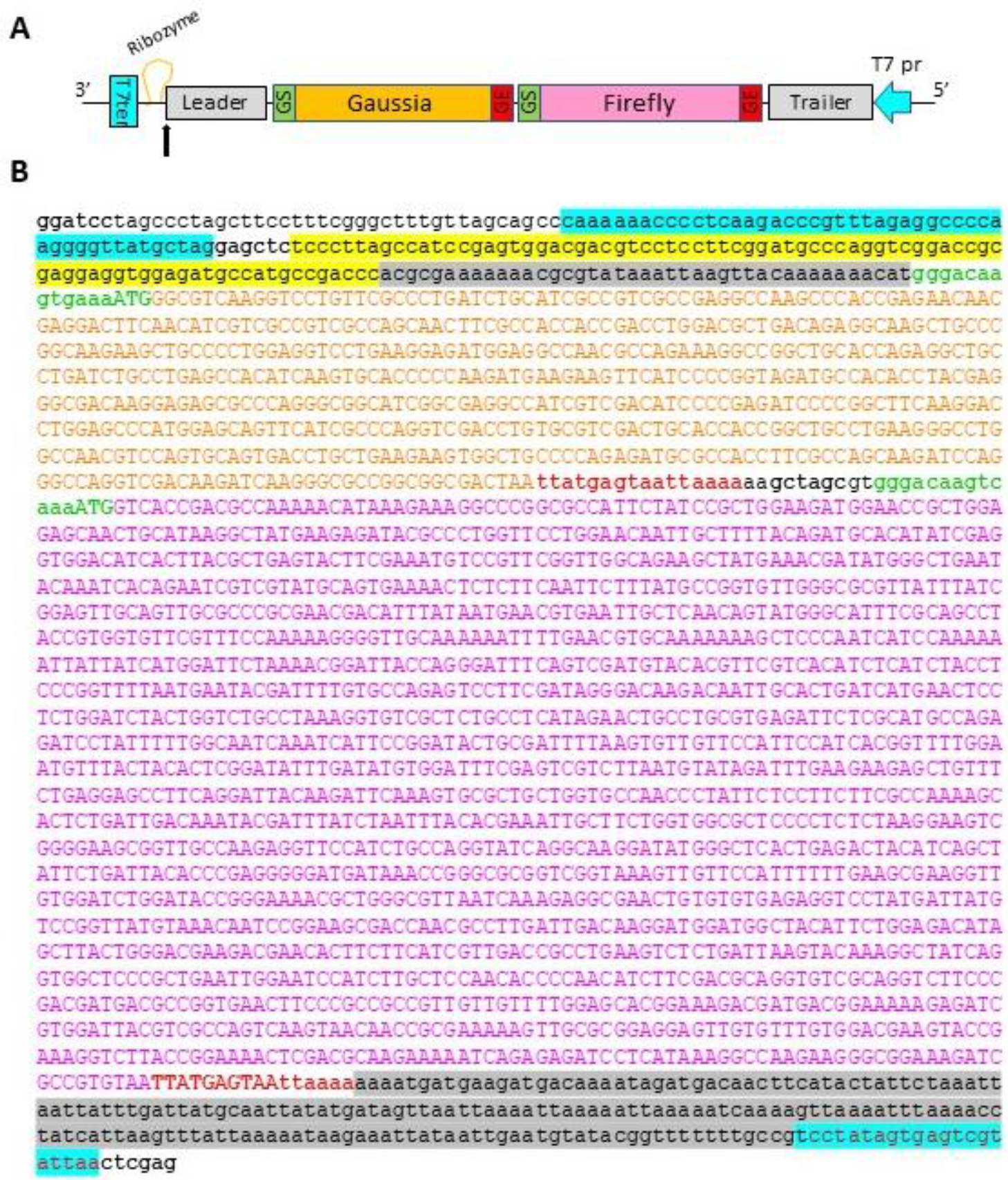
Minigenome construct. A/ Schematic representation of the elements constituting the minigenome sequence. The sequences of the leader and trailer (grey), gene start (GS, green) and gene end (GE, red) are derived from HMPV stain CAN98-87. The full sequence is framed by T7 promoter (T7pr, blue arrow) and terminator (T7ter, blue) sequences. The black arrow indicates the cleavage site induced by the presence of the Ribozyme (yellow). B/ Nucleotide sequence of the HMPV minigenome. The elements are highlighted according to the color code used in A/. Sequences of Firefly and Gaussia luciferase are indicated by uppercase letters. The enzymatic restriction sites are indicated in bold.

**Supplementary Table 1.** Cluster statistics of HADDOCK refined models obtained for HMPV P6/N_NTD_ complex

|  | Cluster 1 | Cluster 2 | Cluster 3 |
| --- | --- | --- | --- |
| HADDOCK score (all) | -47.3 +/- 4.3 | -52.5 +/- 7.9 | -45.5 +/- 3.4 |
| HADDOCK score (top10<sup>a</sup>) | -57.8 +/- 2.8 | -69.3 +/- 4.0 | -49.9 +/- 2.8 |
| Cluster size | 198 | 134 | 36 |
| Cluster RMSD (Å) | 0.95 +/- 0.28 | 0.99 +/- 0.29 | 1.12 +/- 0.56 |
| RMSD from the overall lowest-energy structure Å | 1.69 +/- 0.18 | 0.99 +/- 0.29 | 1.47 +/- 0.24 |
| Van der Waals energy | -15.8 +/- 3.2 | -19.3 +/- 3.4 | -15.8 +/- 3.0 |
| Electrostatic energy | -83.5 +/- 26.1 | -87.1 +/- 20.0 | -77.1 +/- 12.3 |
| Buried Surface Area (Å<sup>2</sup>) | 620.4 +/- 101.1 | 673.7 +/- 102.8 | 606.1 +/- 104.4 |
| *Ionic/polar interactions frequencies* |  |  |  |
| (N)R132--(P)M294 salt-bridge | 2.0 % | 76.9 % | 2.8 % |
| (N)R151--(P)M294 salt-bridge | 75.3 % | 21.6 % | 75.0 % |
| (N)R132-N<sub>η</sub>1/2--(P)I293-O H-bond | 0 % | - | 0 % |
| (N)R151-N<sub>η</sub>1/2--(P)I293-O H-bond | - | 57.5 % | - |
| *# Hydrophobic inter-molecular contacts<sup>b</sup>* |  |  |  |
| (P)L292 | 1.0 +/- 1.2 | 1.0 +/- 1.4 | 0.0 +/- 0.2 |
| (P)I293 | 0.3 +/- 0.8 | 0.3 +/- 0.6 | 0.4 +/- 0.8 |
| (P)M294 | 3.6 +/- 2.3 | 4.2 +/- 2.2 | 3.3 +/- 1.7 |

<sup>a</sup> 10 lowest-energy structure of the cluster ; <sup>b</sup> average number of contacts between Carbon or Sulfur atoms within 3.9 Å

**Supplementary Figure 2.**
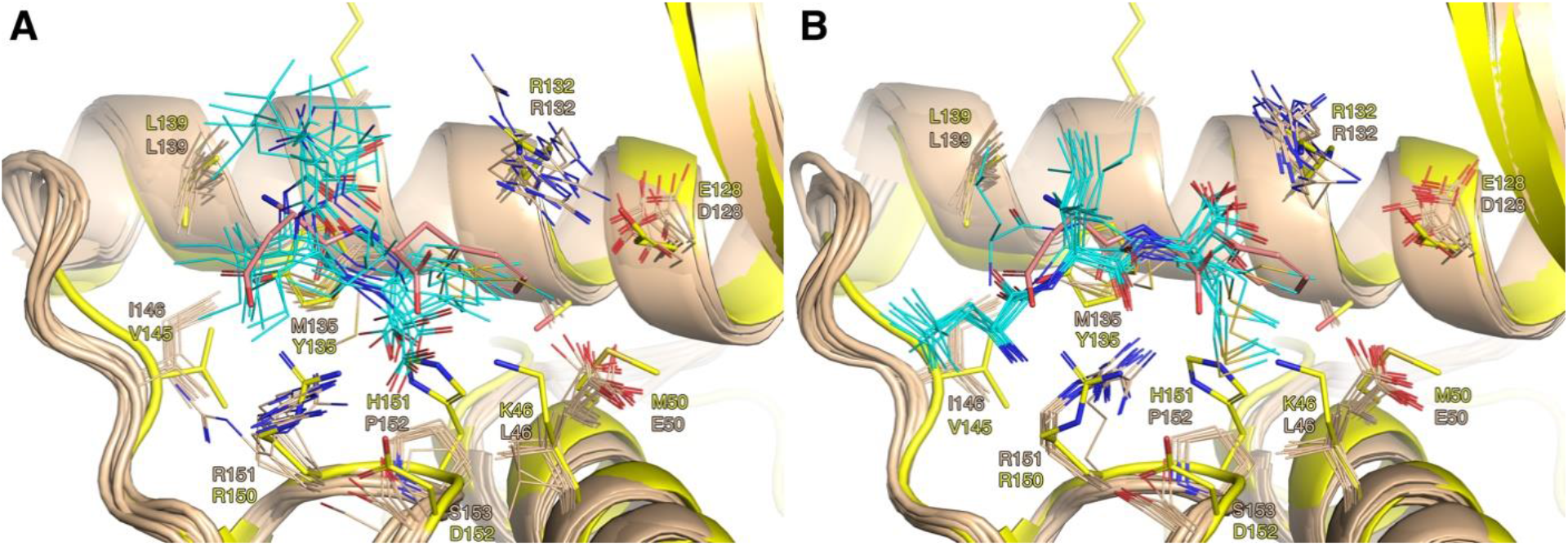
10 best-scoring structures of clusters 1 (A) and 2 (B) from HADDOCK refinement of HMPV P3 (residues L292-I293-M294 of P)/N_NTD_ complex superimposed with RSV P2/N_NTD_. N_NTD_ is colored in beige (HMPV) or yellow (RSV), and HMPV P3 peptide in cyan (HMPV) or RSV P2 peptide (corresponding to residues D240-F241 of P) in pink. Side-chains of residues in contact with the P peptide in HADDOCK models are also shown, along with their RSV equivalent.

